# Resolution and reconciliation of non-binary gene trees with transfers, duplications and losses

**DOI:** 10.1101/064675

**Authors:** Edwin Jacox, Mathias Weller, Eric Tannier, Celine Scornavacca

## Abstract

**Motivation:** Gene trees reconstructed from sequence alignments contain poorly supported branches when the phylogenetic signal in the sequences is weak. When a species tree is available, the signal of gains and losses of genes can be used to correctly resolve the unsupported parts of the gene history. Unfortunately, finding the best (i.e. most parsimonious) resolution has been shown to be NP-hard if transfers are considered as possible gene scale events, in addition to gene originations, duplications and losses.

**Results:** We propose an exact, parameterized algorithm solving this problem in single-exponential time, where the parameter is the number of connected branches of the gene tree that show low support from the sequence alignment or, equivalently, the maximum number of children of any node of the gene tree once the low-support branches have been collapsed. We propose a way to choose among optimal solutions based on the available information. We show the usability of this principle on several simulated and biological data sets. The results show a comparable or better quality than several other tested methods having similar goals, but with a lower running time and a guarantee on the optimality of the solution.

**Availability:** Our algorithm has been integrated into the ecceTERA phylogeny package, available at http://mbb.univ-montp2.fr/MBB/download_sources/16_ecceTERA and which can be run online at http://mbb.univ-montp2.fr/MBB/subsection/softExec.php?soft=eccetera.

## 1 Introduction

Constructing good gene trees is both crucial and very challenging for molecular evolutionary studies. The most common way to proceed is to compute a multiple alignment of nucleotide or protein sequences from a gene family, and search for an evolutionary tree that is most likely to produce this alignment (under some evolutionary model). It is strongly advised to additionally compute statistical supports on the branches of the output tree, as this tells whether they are inferred from the signal of mutations contained in the alignment or are chosen at random in the absence of signal (Felsenstein, 2004). It is very rare that the gene sequences contain enough mutations, but not too many, to support all the branches of a gene phylogeny (Mossel and Steel, 2005). In consequence, it is very rare that a maximum-likelihood tree computed from a multiple alignment reflects the true history of the genes.

A way to approach this true history is to use the information contained in a species tree to correct the branches of a gene tree that are not supported by the alignment. The interpretation of a gene tree in the view of a species tree necessitates the introduction of gene scale events, as the birth of the gene, its death, its replication and diversification by speciation, duplication, and horizontal gene transfer (Szöllősi *et al.,* 2015). If costs are assigned to each gene scale event, each such complete history, called a *reconciliation,* has a total cost. Binary gene trees reconstructed with the additional information of this reconciliation score show a higher quality, according to all tests on methods that are able to perform such a construction: MowgliNNI (Nguyen *et al.,* 2013), ALE (Szöllősi *etal.,*2013), Tera (Scornavacca *et al.,* 2015), TreefixDTL (Bansal *et al.,* 2014) and JPrIME-DLTRS (Sjöstrand *et al.,* 2014) (we enumerate only the methods allowing horizontal gene transfers). However, these methods generally require intensive computing time.

Here, we provide an algorithm that, given a species tree *S* and a gene tree *G* with supports on its branches, computes a modified gene tree *G'* such that all well-supported branches of *G* are in *G'* and, no thusly modified gene tree has a lower reconciliation cost than *G'*. "Well supported" is defined by a threshold chosen by the user, or by an adaptive method which sets this support according to the algorithm complexity. This problem reduces to the reconciliation of non-binary gene trees with binary species trees, which has been showed to be NP-complete (Kordi and Bansal, 2015). We employ amalgamation principles (Scornavacca *et al.,* 2015) to prove its fixed-parameter tractability with respect to a reasonable parameter: the maximum number of children of any node of the gene tree once the low-support branches have been collapsed. We provide an implementation and we propose a way to choose among all optimal solutions according to the (both low and high) supports of the branches of the input tree.

We show on both simulated and real data sets that our method produces gene trees of comparable quality than those constructed by competitive methods, often in a smaller amount of time. The software is easy to use, so that anyone having a gene tree with supports and a species tree can quickly obtain a better-quality gene tree, or even correct a whole database in reasonable time.

## 2 An FPT algorithm for reconciling gene trees with polytomies

### 2.1 Reconciliation of binary gene trees

For our purposes a *rooted phylogenetic tree T*=(*V*(*T*), *E*(*T*), *r*(*T*)) is a tree, where *V*(*T*) is the set of nodes, *E*(*T*) is the set of arcs, all oriented away from *r*(*T*), the root. For an arc (*x, y*) of *T*, we call *x* the *parent* of *y*, and *y a child* of *x*. The number of children of *x*, denoted by *k_x_*, is called the *out-degree* of *x*. If a path from the root to *y* contains *x*, then *x* is an *ancestor* of *y* and *y is a descendant* of *x*. This defines a partial order denoted by *y≤T x*, and *y<T x* if *x ≠ y*. The subtree of *T* that is *rooted at* a node *u* of *T* (denoted by *T_u_*) is the result of deleting all nodes *v* with *v≤T u* from *T*. Nodes with no children are *leaves*, all others are *internal nodes*. The set of leaves of a tree *T* is denoted by *L*(*T*). The leaves of *T* are bijectively labeled by a set *ℒ*(*T*) of labels. A tree is *binary* if all internal nodes have exactly two children. A tree *T* is *dated* if a total order *≤θ* on internal nodes is given that extends *≤T*. Each internal node then implies a *date* between 1 and |*V*(*T*) | —|*L*(*T*)| (the root) and all the leaves are assumed to have date 0. The *dated subdivision T'* of a dated tree *T* is obtained by replacing each arc (*x, y*) by a path containing *k* additional nodes, where *k*+1 is the difference between the date of *x* and the date of *y*.

A reconciliation involves two rooted phylogenetic trees, a *gene tree G* and a *species tree S*, both binary. Their relation is set by a function *s*: *ℒ*(*G*)*→ℒ*(*S*), which means that each extant gene belongs to an extant species. Note that *s* does not have to be injective (several genes of *G* can belong to the same species) or surjective (some species may not contain any gene of *G*). A *reconciliation α of G* in *S* is a mapping of each internal node *u* of *G* into a sequence (*α*(*u*)_1_, *α*(*u*)_2_, …) of nodes of *S* if *S* is undated or nodes of the dated subdivision *S'* of *S* if *S* is dated. Herein, for each *i* ≥ 1, we have *α*(*u*)*_i+1_* ≤ *T α*(*u*)*_i_* and *α*(*u*)*_i_* satisfies the constraints of any one of the possible events among duplication (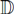), transfer (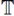), loss (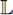), or speciation (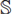) - see Appendix 1 for details. This ensures that a coherent gene history can be extracted from α. Given costs for individual 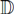, 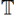 and 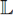 events (it is usually assumed that speciation does not incur cost), denoted *δ, τ* and *λ* respectively, a reconciliation *α* is assigned the cost *c(α):= dδ+tτ+lλ*, where *d, t* and *l* denote the respective numbers of events of type 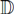, 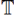 and 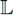 implied by *α*. We denote by *ℛ(G, S)* the set of all possible reconciliations of *G* in *S* and define 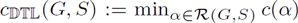, that is, the minimum cost over all possible reconciliations of *G* in *S*. We call 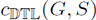 the *cost* of *G* with respect to a species tree *S*.

#### Problem 1.

Most Parsimonious Reconciliation *(MPR)*

**Instance:** *a (dated) binary species tree S, a binary gene tree G, costs δ, τ, λ for respective 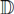, 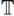, 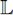 events*

**Output:***a reconciliation of minimal cost of G in S*

This problem can be solved in *O*(|*S*|*^2^* · |*G*|) time (Doyon *et al.,* 2010) for dated trees, and *O*(|*S*| · |*G*|) time for undated ones (Bansal *et al.,* 2012). In the following, we turn our attention to non-binary gene trees and we consider the species tree as dated. Nevertheless, every result is also valid for the undated case with better complexity, see Section 3.3.

### 2.2 Resolution of non-binary gene trees

If a node in a tree *T* has more than two children, we call this node a *polytomy*. Note that a node *α* of *T* partitions *L*(*T*) into two sets, the descendants of *α* and all others. Given a gene tree *G* with at least one polytomy, a binary tree *G'* is called *binary resolution* of *G* if *G* can be obtained from *G'* by contracting edges. We denote by BR(*G*) the set of all binary resolutions of *G*.

#### Problem 2.

Polytomy Solver under the 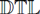 framework

**Instance:** *a dated binary species tree S, a gene tree G, costs δ, τ, λ respectively for 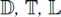 events*

**Output:** *a binary resolution G' of G minimizing* 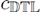 (*G'*, *S*)

This problem is known to be NP-hard (Kordi and Bansal, 2015), and can be solved in polynomial time for *τ=∞* (Lafond *et al.,* 2012). A brute-force approach would need to generate all binary resolutions. There are 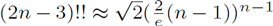 different rooted binary trees on n leaves, which gives 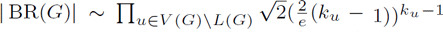 different binary resolutions of *G*. This yields an algorithm with time complexity *O*(|*S*|^2^ · |*G*| · |*BR*(*G*)|). Using the following result, we can improve on the brute-force approach.

#### Theorem 2.1.

*Let S be a species tree, let G be a gene tree with a unique polytomy at its root. Let G', G^*^ G ∈ BR(G) and let u', u* be vertices of G' and G^*^, respectively, such that ℒ(G'_u'_), =ℒ(G^*^_u^*^_. Let α ∈ ℛ(G', S) and β ∈ ℛ(G^*^, S) such that α(u')_1_=β(u^*^)_1_. Then, the result of replacing the restriction of α to G'_u'_, by the restriction of β to G^*^_u^*^_ is a reconciliation of ℛ(G', S)*.

We do not give a proof of Theorem 2.1 here because it will be a consequence of our main Theorem 2.2. Nevertheless, we note here that the dynamic programming algorithm by Doyon *et al.* (2010) with the help of Theorem 2.1 permits us to solve Problem 2 with a lower complexity: Whenever we encounter a polytomy *u* in its bottom-up approach, we store, for each *v* ∈ *V*(*S*), the minimum cost of a reconciliation associating *u* with *v* over all binary resolutions of *G_u_*.

#### Observation 2.1.

*A solution for Problem 2 can be found by solving polytomies one by one in a bottom-up approach, with a time complexity of 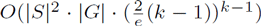, where k is the maximum number of children of any node in G*.

While Observation 2.1 already implies that Problem 2 is fixed-parameter tractable with respect to the maximum out-degree in *G*, it remains interesting to search for a single-exponential-time algorithm. The idea for such an algorithm comes from the amalgamation principle.

### 2.3 Amalgamation of gene trees

A node *u* of a binary tree *T* is said to generate the *clade C*(*u*)=*ℒ*(*T_u_*). If *u* has distinct children *u_r_* and *u_l_*, we also say that *u* generates the tripartition (*C*(*u*), {*C*(*u*_l_), *C*(*u_r_*)}), otherwise, *u* generates the *trivial* tripartition (*C*(*u*), {*Ø, Ø*}). Herein, we call *C*(*u*) the *domain* of the tripartition. A binary tree *T* generates a set of clades and tripartitions, respectively denoted 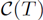 and Π(*T*), which are the clades and tripartitions generated by its nodes.

More generally, for a set of labels *L*, a clade is a subset of *L* and a tripartition is a tuple (*C*, {*C_r_*, *C_l_*}) such that the clades *C_r_, C_l_* partition the clade *C*. Let Π be a set of tripartitions on *L*. We denote the set of clades present in Π by 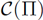. Further, Π is said to be *complete* if it contains *L* as the domain of some tripartition, and for each tripartition (*A*, {*B, C*}) either *B=C=Ø* or Ø contains tripartitions with *B* and *C* as respective domains. It is easy to see that the set of tripartitions generated by a binary tree is always complete, and conversely, for any complete set of tripartitions Π on *L*, there is at least one binary tree *G* with *ℒ*(*G*)=*L*, which generates a subset of Π. The amalgamation problem is to generate one minimizing the reconciliation cost with respect to a species tree:

#### Problem 3.

Amalgamation under the 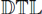 framework

**Input:** *a complete set of tripartitions Π on a set of labels L, a dated binary species tree S, a labeling function s:L → ℒ(S) and costsδ, τ, λ respectively for 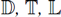 events*

**Output:** *a binary gene tree G minimizing 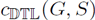 over all binary gene trees G' with ℒ(G')=L and such that the set of tripartitions generated by G' is a subset of Π*

This problem can be solved in *O*(|*S*|^2^ · |Π|)time (Scornavacca *et al.,* 2015). Given a non-binary tree *G*, we can generate a complete set of tripartitions containing all tripartitions of any binary resolution of *G* as follows: for each node *u* of *G* with child set {*u_1_, …, u_t_*}, and for each *I ⊆* {*u_1_, …, u_t_*}, *u* generates a tripartition (*A*, {*B, C*}) such that *B ⋃ C=A ⊆ ⋃*_i∈I_*C*(*u_i_*) and none of *B* or *C* overlap any *C*(*u_i_*)(overlapping means containing some elements but not all). The union of all these tripartitions is the set of tripartitions of *G*, noted Π(*G*) as in the binary case.

Thus, the solution proposed in (Scornavacca *et al.,* 2015) yields a novel algorithm for the resolution of polytomies.

#### Theorem 2.2.

*Let G be a non-binary gene tree. An amalgamation solution on Π(G) is an optimal binary resolution of G*.

The complexity of this algorithm is *O*(|*S*|^2^ · |Π(*G*)|). We can bound the size of Π(*G*) using the following statement.

#### Lemma 2.3.

*For any gene tree G, |Π(G)|=O(|G| · (3^k^ — 2^k+1^)), where k is the maximum out-degree of any node in G*.

#### Proof.

Let *u* be a node of *G*, and *u_1_, …, u_t_* its children. Recall that, for each *I ⊆ {u_1_, …, u_t_}*, *u* generates a tripartition (*A*, {*B, C*}) such that *B⋃C=A ⊆ ⋃*_*i∈I*_*C*(*u_i_*) and none of *B* or *C* overlap any *C*(*u_i_*). Any tripartition (*A*, {*B, C*}) generated by *u* is isomorphic to a partition of {*C*(*u_1_*), …, *C*(*u_t_*)} into three sets, *B, C,* and *⋃*_*i*_*C*(*u_i_*)\*A* with *B* ≠ Ø and C ≠ Ø. The number of partitions of *t* elements into three sets is 3*^t^* and 2 · 2*^t^* of them have B=Ø or C=Ø. Avoiding to count the (unique) partition with B=C=Ø twice, we get 3*^t^* — 2^t+1^+1 such partitions. Finally, we can remove half of the remaining partitions by symmetry of *B* and *C* and arrive at a count of 1/2(3*^t^* — 2*^t+1^*+1). Thus, there are 1/2(3*^t^* — 2*^t+1^*+1) such partitions and, hence, this is also an upper bound on the number of tripartitions generated by *u*. Summing over all vertices of *G*, the total number of tripartitions generated by *G* is then bounded by |*G*|· 1/2(3*^k^* — 2*^k^+^1^* +1) where *k* is the maximum out-degree in *G*.

This leads to the main theoretical result of the paper

#### Proposition 2.4.

*For any gene tree G, Problem 2 can be solved in O(|S|^2^ · |G| · (3^k^ — 2^k+1^)) time for dated species trees, where k is the maximum out-degree in G*.

## 3 Practical issues

In order to turn the algorithmic principle described in the previous section into a workable method for biological datasets, we have to handle three issues: one is that the position of the root in the gene tree is usually unknown; a second is that species trees are usually undated; the last and most difficult one is the choice between multiple solutions. Indeed in some cases the solution space of the problems defined in the previous sections is huge and two different solutions can be far apart. But some information from branch supports - encoded in the tripartitions to be used in amalgamation algorithm - can be used to efficiently find a good solution. We address this issue first.

### 3.1 Scoring tripartitions as a guide in the solution space

Given a multiset of tripartions Π on *L*, the *conditional probability* of a tripartition π=(*C_1_*, {*C_2_, C_3_*}) in Π is defined as the ratio *f_Π_(π)/f_Π_(C_1_)*, where, for each clade and tripartition in Π, *f_Π_*(·) denotes its frequency in Π (Höhna and Drummond, 2012). The *conditional clade probability* of a binary tree *G* such that *ℒ*(*G*)=*L*, denoted by 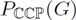, is defined as the product of the conditional probabilities of all tripartitions in Π(*G*).

We propose to output, among all solutions to the non-binary reconciliation problem, one that maximizes the conditional clade probability. For this we construct an artificial probability space, where the conditional probability of a tripartition π=(*C_1_*, {*C_2_, C_3_*}) is still *f_Π_*(π)/*f_Π_*(*C_1_*), but where *f* is redefined using information from branch supports (low or high) of the input tree. The rationale is that we use the support of a branch to approximate the frequency of the corresponding clade in an imaginary sample G of binary resolutions, while other branches not present in the input tree are equiprobable. So we have to assign a probability to each clade and, in the following, we explain precisely how.

*Frequency of clades*. Let *G^B^* be a rooted binary tree with supports on its branches and let *G* be the multifurcated gene tree obtained from *G^B^* by contracting unsupported branches (according to a given threshold), we define a support *f*(*C*_1_) for each clade *C_1_* generated by *G*. If 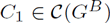, then *f*(*C_1_*) is its support, i.e. the support of the branch leading to the clade. ^1^Otherwise, there is a clade in *G^B^* that is incompatible with *C_1_*. Among all such clades, let *C_0_* be one that maximizes *f*(*C*_0_). Then, we use the knowledge of the frequency of *C_0_* to infer the frequency of *C_1_* in our imaginary sample 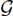 by assuming that the ratio of trees generating *C_1_* to trees not generating *C_0_* is the same in 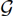 as in *BR*(*G*). Thus, the support of *C_1_* is defined as 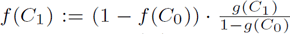, where *g*(*C*) is the frequency of a clade *C* in in *BR*(*G*). To compute *g*(*C*), suppose *C* is generated by a vertex *u* of *G* with *n*(*u*) children, and that *n*(*C*) is the number of children of *u* “contained” in *C*. Then:

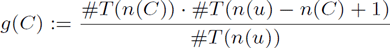

where *#T*(*k*) is the number of rooted binary trees with *k* leaves, i.e. (2*k* — 3)!!.

For example, given the trees in Figure 1, the support of the clade {*C, D*} is 0.5 (i.e. 50/100), while the support of {*A, B, C*} (which conflicts with {*C, D*}) is 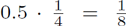 (note that 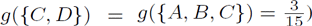).

**Fig. 1.**
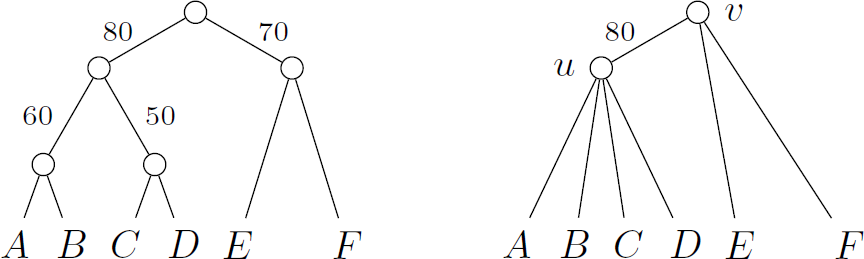
A binary gene tree *G^B^* (left) and a non-binary one (right) obtained from *G^B^* by suppressing edges with a bootstrap lower than 80.

*Frequency of tripartitions*. Let π=(*C^1^*, {*C_2_, C_3_*}) be a tripartition. Let *BR*(*G*)_1_ be the set of binary resolutions of *G* that generate *C_1_* and let 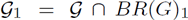 be the part of our imaginary sample whose trees generate *C_1_*. If π ∈ Π(*G^B^*), we define *f*(π):=min(*f*(*C_i_*), *f*(*C_2_*), *f*(*C_3_*)). Otherwise, either *G^B^* generates *C_1_* and a tripartition Π' on the same domain *C_1_* as π, or *G^B^* does not generate *C_1_*.

In the first case, we use the knowledge of the frequency of π' and *C_1_* to infer the frequency of π in our imaginary sample by assuming that the ratio of trees generating π to trees not generating π' is the same in 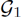 as in *BR*(*G*)_1_. Thus, *f*(π):=(*f*(*C_1_*)−*f*(π'))· *ĝ*(π)/(*g*(*C_1_*)−*ĝ*(π')), where *ĝ*(n) is the frequency of n in BR(G). In the second case, we assume that the frequency of trees generating π is the same in 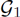 as in *BR*(*G*)_i_. Thus, *f*(*π*):=*f*(*C_1_*) · *ĝ*(π)/*ĝ*(*C_1_*), where *ĝ*(π) is the frequency of π in *BR*(*G*).

The frequencies *ĝ* can be computed as follows. If π=(*C_1_*, {*C_2_*, *C_3_*}) is generated by *u*, then we denote by *n*(*C_i_*) the number of children of *u* whose clades are contained in *C_i_*. We define *ĝ*(*π*) as

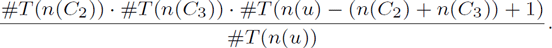

For example, given the trees in Figure 1, we have the following:

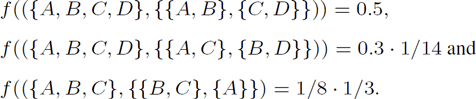

Finally, conditional clade probability and reconciliation cost can be combined by weighting their ratio (see Scornavacca *et al.* (2015)).

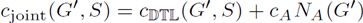

where G' is a gene tree in BR(G) and the parameter *c_A_* weights the contribution of the “sequence contribution” *N_A_*(*G*) to the cost, defined as

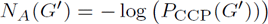

We will use such a joint score in the applications.

### 3.2 Unrooted gene trees

Phylogenetic trees are always rooted, but often the position of the root is unknown. The method in the previous section can be used on an unrooted gene tree *G* to account for the uncertainty on the position of the root, without additional complexity as follows. First, we call *G_r_* the rooted tree obtained by rooting *G* arbitrarily on an internal edge *e*, and 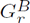 the rooted version of *G^B^*, also rooted on *e*. Then we compute the set Π(*G_r_*) of tripartitions as defined in Section 2.3. To obtain Π(*G*), we also consider, for each non-trivial tripartition (*C_1_*, {*C_2_, C_3_*}) ∈ Π(*G_r_*), the two other possible tripartitions on *ℒ*(*G*) that are implied by a different placement of the root, namely ((*ℒ*(*G*) \ *C_1_*) ⋃ *C*_2_, {*ℒ*(*G*)\*C*_1_, *C_2_*}) and ((*ℒ*(*G*) \ *C*_1_) ⋃ *C*_3_, {*ℒ*(*G*) \ *C*_1_, *C*_3_}). To these tripartitions, we add the set Π':={(*ℒ*(*G*), {{*l*}, *ℒ*(*G*) \ {*l*}})|*l*∈ *ℒ*(*G*)} of all trivial tripartitions.

Each edge *e*=(*u, v*) of the rooted binary tree 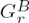 *induces* two clades *C_u_*(*e*) and *C_υ_*(*e*), which correspond to the label sets of the leaves of the two subtrees created by removing *e*. We say that *C_u_*(*e*) is generated by *u* and *C_υ_*(*e*) by *υ*, and we associate to them the support of *e*. Then, we redefine the set 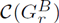 as the set of clades induced by all edges of 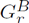. Given the thusly defined set 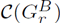, the support of each clade of 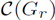 is then computed as in the rooted case.

Finally, we describe how to give a support value to each tripartition of *G* in the unrooted case: The support of each tripartition π ∈ Π(*G_r_*) is computed w.r.t. 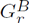 as described in the previous section, and the support of the two other possible tripartitions on *ℒ*(*G*) that are implied by a different placement of the root is the same as the support of π. All tripartitions in Π' have support equal to 1.

### 3.3 An undated variant

The method discussed by Scornavacca *et al.* (2015) has been conceived for dated binary species trees, but can easily be adapted to undated ones, while respecting all previously mentioned results, with a slight correction concerning the complexity.

Indeed, reconciliations for undated species trees can be computed in *O*(|*S*| · |*G*|) time with an algorithm described by Bansal *et al.* (2012). Adapting this algorithm to the amalgamation framework can be done by the same technique used to adapt the *O*(|*S*| · |*G*|) time algorithm of Doyon *et al.* (2010) for dated species trees reconciliation to the amalgamation framework as done in (Jacox *et al.,* 2016). Thus, our result translates to an *O*(|*S*| · |*G*| · |Π|)-time algorithm-by Lemma 2.3, an *O*(|*S*| · |*G*| · (3*^k^*–2*^k^+^1^*))-time algorithm - for undated species trees.

### 3.4 Adaptive compromise between the amount of correction and the computational complexity

The threshold for deciding if a branch is well-supported or not is, in principle, user-defined. However, in the experiments, we required that, in the multifurcated tree resulting from the contraction of unsupported branches, an internal node has at most 12 children. This is to avoid the combinatorial explosion and to keep the method fast. So we adopted a strategy of increasing the threshold by 0.02 steps until the 12 maximum children property was satisfied.

## 4 Application on simulated and biological data

In this section, we use three different data sets, two simulated and one from microbial genomes, to compare the performance of our algorithm (called ecceTERA in the following) with six different gene tree reconstruction methods: TERA (Scornavacca *et al.,* 2015), ALE (Szöllősi *etal.,* 2013), TreeFix-DTL (Bansal *et al.,* 2014), MowgliNNI (Nguyen *et al.,* 2013), JPrIME-DLTRS (Sjöstrand *et al.,* 2014) and PhyML (Guindon *et al.,* 2010). The five first use information from the sequences and the species tree, while the last uses only information from the sequences.

### 4.1 Simulated Proteobacteria Data Set

The proteobacteria data set is the one constructed to test MowgliNNI and made available by Nguyen *et al.* (2013). Starting with a dated phylogeny of 37 proteobacteria (David and Alm, 2011):

- 1000 evolutionary histories comprising D, T, and L events were simulated along the species tree according to a birth and death process;
- sequences were simulated along these true gene trees under the GTR model using Seq-Gen (Rambaut and Grass, 1997);
- RaxML (Stamatakis, 2014) was used to estimate gene trees (along with 100 bootstrap trees) from these sequences under the GTR model.

We refer to the section “Simulated gene trees and evolutionary histories” of Nguyen *et al.* (2013) for more details on how the data set was composed. Some of the test results are taken from the same procedure proposed by Scornavacca *et al.* (2015).

We ran TreeFix-DTL with default parameters, GTR with a gamma distribution as models of evolution, and as a starting tree the RaxML tree. The RaxML tree (with bootstrap values) was also the input given to MowgliNNI and ecceTERA that were run with a threshold for weak edges equal to 70. As in Scornavacca *et al.* (2015), JPrIME-DLTRS was run on the sequence alignments with JTT with a gamma distribution as model of evolution, 100 000 iterations, a thinning factor of 10 and a time out of 10h. Finally, TERA and ALE were run on the set of bootstrap trees with default parameters, except for the weight of amalgamation *cA*, fixed to 0.1 for TERA as for ecceTERA.

The accuracy - defined as normalized Robinson-Foulds distance to the true tree - of TERA, ecceTERA and ALE is comparable and higher than the one of JPrIME-DLTRS and RaxML (see Figure 2). The method that outperforms the others in this case is TreeFix-DTL. This is probably due to the fact that the data set was simulated with the same model used to estimate the gene trees with TreeFix-DTL. Note also that TreeFix-DTL is slower than ecceTERA, with an average runtime of 37 minutes versus 15 seconds.

**Fig. 2.**
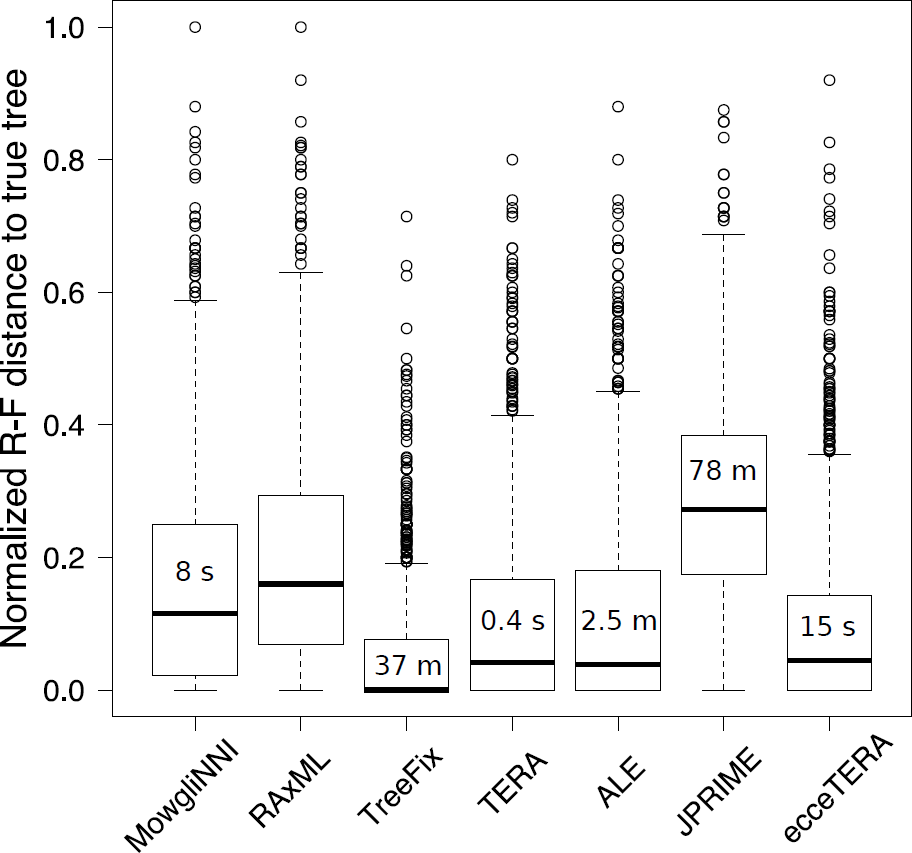
Accuracy of several methods on the simulated proteobacteria data set: we measure the Robinson-Foulds distance of the reconstructed tree to the true tree for all 1000 gene trees, for 7 methods. Computing times - except for the computation of the RAxML trees - are given in the boxes (s=seconds, m=minutes). In terms of quality, TreeFix-DTL achieves the best accuracy but is relatively slow. ecceTERA is comparable with the second best methods, both in accuracy and computing time.

### 4.2 Cyanobacteria Data sets

The biological and simulated cyanobacteria data sets used here have been made public by Szöllősi *et al.* (2013) at http://datadryad.org/resource/doi:10.5061/dryad.pv6df.

Their construction consisted in selecting 1099 gene families from 36 cyanobacteria species, related by a known dated species tree. These families were retrieved from HOGENOM (Penel *et al.,* 2009), and were selected for their reasonable size and representativity. To obtain the biological data set, multiple-alignments on these families were computed with Muscle (Edgar, 2004).

To obtain the simulated data set, from each multiple-alignment of the real data set, a sample of 1000 trees was computed with Phylobayes (Lartillot *et al.,* 2009), and an amalgamated tree was reconstructed with ALE (Szöllősi *et al.,* 2013). This tree was used to simulate multiple-alignments of artificial sequences evolved along this tree under an LG model with a gamma distribution. This multiple-alignment is the input of our simulated data set. See (Szöllősi *et al.,* 2013) for more detail on how the data sets were generated.

*Tests on the simulated data set.* A tree was computed for each simulated multiple-alignment using PhyML (Guindon *et al.,* 2010), with an LG+Γ4+I model and SH branch supports.

For this data set, the accuracy of TERA, ALE, TreeFix-DTL, MowgliNNI, JPrIME-DLTRS and PhyML were compared in (Scornavacca *et al.,* 2015, Figure 2(a)), see Section 2.5 of the same paper for more details on the input/parameters used to generate the results. In Figure 3, we compare these results with those of ecceTERA, again on unrooted gene trees and dated species tree, for three different thresholds of weakly supported edges (0.8, 0.7 and 0.6). We report the results of ALE and PhyML from Scornavacca *et al.* (2015), plus the results for MowgliNNI for the same thresholds used for ecceTERA. Both ecceTERA and Mowgli were given the “simulated” PhyML trees with SH branch supports. Finally, we report the accuracy of ALE when using as input 100 sample trees (ALE 100 in Figure 3) among the 10k trees provided, mimicking the information contained in a set of bootstrap trees.

**Fig. 3:**
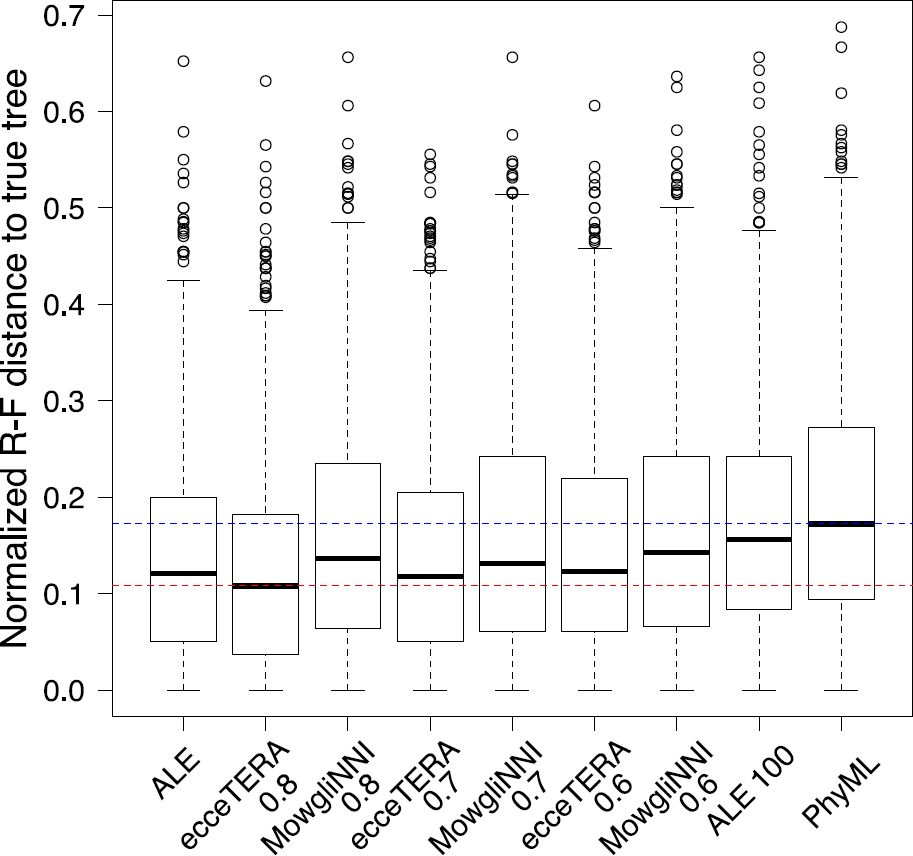
Accuracy of several methods on the simulated cyanobacteria data set: ecceTERA with a threshold of 0.8 achieves better accuracy than all other methods.

Figure 3 shows that ecceTERA with athreshold of 0.8 achieves abetter accuracy than all other methods. Moreover, the accuracy of ecceTERA increases with the threshold for weak edges, while this is not the case for MowgliNNI. It is also worth noting that the accuracy of ALE decreases considerably when used on small samples of trees. These results are even more interesting when considering that ecceTERA is the fastest method of the bundle (see Table 1 and Figure 2(a) of (Scornavacca *et al.,* 2015)). Although ecceTERA has similar running times as TERA, it requires much less time to construct the input (as its input is the PhyML tree with SH branch supports). Thus, the ecceTERA strategy is the fastest (considering computation+input preparation time) among all compared strategies.

*Tests on the biological data set*. From each multiple alignment, a maximum-likelihood tree was computed with PhyML (Guindon *et al.,* 2010) with SH branch supports. These PhyML trees were corrected using ecceTERA with a threshold for weak edges equal to 0.8. The weight of amalgamation *cA* was estimated, with starting value 1. Gene trees were considered as unrooted and the species tree as dated.

We measured the quality of the corrected trees compared to that of the maximum-likelihood trees in two ways. First, we compared the likelihoods according to the multiple-alignments. Of course, the PhyML trees always have a better likelihood because they are optimized with respect to this criterion. But it is interesting to note that for 80% of the trees, an Approximate Unbiased (AU) test performed with Consel (Shimodaira and Hasegawa, 2001) did not reject the ecceTERA tree. So, in a vast majority of the cases, the ecceTERA and PhyML trees are equivalent regarding their sequence likelihoods. Second, we compared the two sets of trees with respect to their implications for the evolutionary dynamics of genomes: we counted the number of genes present in ancestral species according to reconciled gene trees, and compared it with the number of genes present in extant species (in our data set which contains a subset of gene families). Ancestral genes were inferred from reconciliations. The results are shown in Figure 4.

**Fig. 4:**
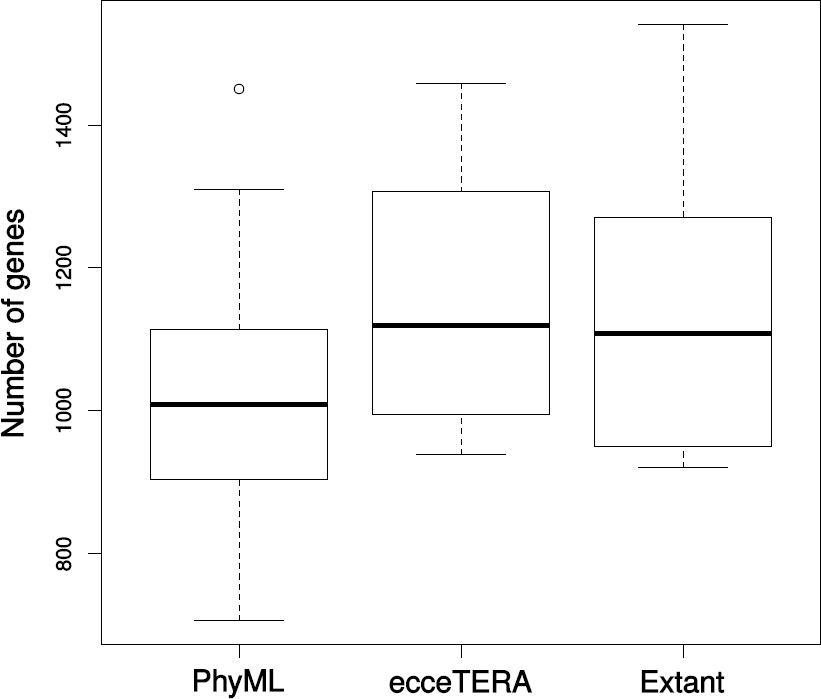
Number of genes for extant species (right), or ancestral species (left and middle boxes), reconstructed with reconciled trees from PhyML (left) and ecceTERA (middle). Better trees yield more plausible genome histories.

We see that the genome histories are much more stable according to ecceTERA trees. According to PhyML trees, ancestral genomes were much smaller than extant genomes, which is not a plausible hypothesis. Indeed there is no argument for why PhyML trees would capture a biological signal that more sophisticated methods constructing statistically equivalent trees from the sequence point of view, would miss. This result argues for the correction step, quickly achieved by ecceTERA, for every evolutionary study.

## 5 Conclusions

Gene trees are a precious resource for biologists. They allow us to annotate genomes, to define species taxonomies, and to understand the evolution of traits, adaptation, and modes of genome evolution. They are also used to reconstruct ancestral genomes and understand the history of relations between organisms and their environment on a long time scale. Thus, reliable gene trees are crucial for many biological results (see for example Groussin *et al.* (2015)).

Standard software constructing gene trees from multiple sequence alignments are widely used. Although they provide better quality gene trees, it is less the case for species tree aware methods. First because a species tree is not always available, and second because of the computational investment most methods require to output a gene tree.

We propose here a method that can be easily used by biologists to quickly correct the output of a gene tree computed from a multiple sequence alignment, provided that branch supports and a species tree are available. The software is built on an FPT algorithm which is derived from recent advances in gene tree amalgamation principles. Its complexity is an exponential function of the maximum degree of the input tree, which reduces to the number of connected branches with low support. What a low support is depends on a threshold chosen by the user. Thus, a compromise between the extent of the correction and the computing time is easily achieved. On all of our data sets, on several dozens of species and several thousand of genes, we arrived quickly at a result that is always significantly better than methods based on multiple sequence alignments only, and whose quality is equivalent to the computationally more intensive integrated methods.

## Acknowledgements

All analyses were performed on the computing cluster of the Montpellier Bioinformatics Biodiversity (MBB) platform.

## Footnotes

1 Note that supports have to be numbers between 0 and 1, and thus bootstrap values should be divided by the size of the bootstrap sample.

## 1 Appendix

Given a binary gene tree *G* and an undated, respectively dated, species tree *S* such that *ℒ*(*G*)⊆ *ℒ*(*S*), an undated, respectively dated, 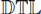 reconciliation *α* is a function that maps each node *u* of *G* onto an ordered sequence of nodes of *S*, respectively of the subdivision *S'*, denoted *α*(*u*)=(α_1_(*u*), *α*_2_(*u*), …, *α*_*ℓ*_(*u*)). Seven basic events are considered by the 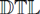 model: speciation (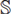), duplication (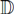), transfer (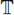), contemporary (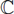), transfer+loss (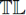) and speciation+loss (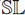) events, plus the additional no event (Ø). 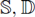 and 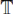 events are self-explanatory. A Ø event denotes that a gene crosses a time boundary, with no other event happening. A contemporary event 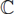 associates a leaf *u* of *G* with a leaf *x* of *S'* such that *s*(*u*)=*s*(*x*). A duplication followed immediately by a loss, i.e. a 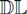 event, can occur an arbitrary number of times in a reconciliation, making the solution space infinite. To prevent this, loss events are never considered alone but are always coupled with either a speciation or a transfer event. Thus, an 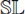 event is a speciation where the gene is lost in one of the two derived species, while a 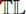 event is a 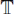 event where the transferred gene is not kept in the descendants of the donor species. Consecutive 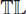 events are not allowed in order to prevent the solution space from being infinite. Note that 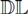 events and consecutive 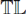 cannot happen in parsimonious reconciliations.

More formally, for each pair of nodes *u* of *G* and *α_i_*(*u*) of *S* (denoting *α*_i_(*u*) by *x* below), *α* is said to be an *undated reconciliation* between *G* and *S* if and only if exactly one of the following events occurs for each pair of nodes *u* of *G* and *α_i_*(*u*) of *S* (denoting *α_i_*(*u*) by *x* below):

a) if *x* is the last node of *α*(*u*), one of the cases below is true:
1. 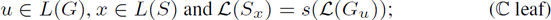
2. 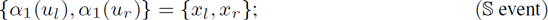
3. 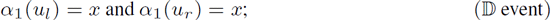
4. *α*_1_(*u_l_*)= *x*, and *α*_1_(*u_r_*) is any species node that is a not a descendant or ancestor of *x*;
or 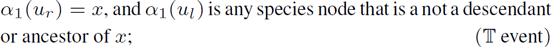
b) otherwise, one of the cases below is true:
5. 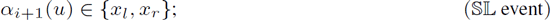
6. 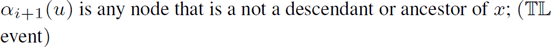

The function a is said to be a *dated reconciliation* between *G* and *S* if and only if exactly one of the following events occurs for each couple of nodes *u* of *G* and *α_i_*(*u*) of *S'* (denoting *α_i_*(*u*) by *x'* below):

a) if *x'* is the last node of *α*(*u*), one of the cases below is true:
1. 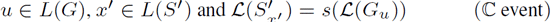
2. 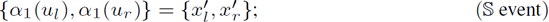
3. 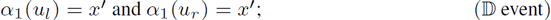
4. *α_1_*(*u_l_*)=*x'*, and *α*_1_(*u_r_*) is any node other than *x'* having height *h*(*x'*)
or 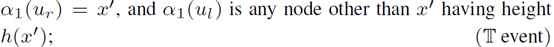
b) otherwise, one of the cases below is true:
5. *x'* is an artificial node and *α_i+1_*(*u*) is its only child; (Ø event)
6. 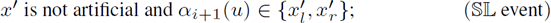
7. 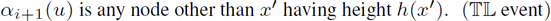

